# Association of predicted deleterious single nucleotide polymorphisms with carcass traits in meat-type chickens

**DOI:** 10.1101/285924

**Authors:** Priscila Anchieta Trevisoli, Gabriel Costa Monteiro Moreira, Clarissa Boschiero, Aline Silva Mello Cesar, Juliana Petrini, Mônica Corrêa Ledur, Gerson Barreto Mourão, Luiz Lehmann Coutinho

## Abstract

In previous studies, we used genome wide association (GWAS) to identify quantitative trait loci (QTL) associated with weight and yield of abdominal fat, drumstick, thigh and breast traits in chickens. However, this methodology assumes that the studied variants are in linkage disequilibrium with the causal mutation and consequently do not identify it. In an attempt to identify causal mutations in candidate genes for carcass traits in broilers, we selected 20 predicted deleterious SNPs within QTLs for association analysis. Additive, dominance and allele substitution effects were tested. From the 20 SNPs analyzed, we identified six SNPs with significant association (p-value <0.05) with carcass traits, and three are highlighted here. The SNP rs736010549 was associated with drumstick weight and yield with significant additive and dominance effects. The SNP rs739508259 was associated with thigh weight and yield, and with significant additive and allele substitution effects. The SNP rs313532967 was associated with breast weight and yield. The three SNPs that were associated with carcass traits (rs736010549, rs739508259 and rs313532967) are respectively located in the coding regions of the *WDR77, VWA8* and *BARL* genes. These genes are involved in biological processes such as steroid hormone signaling pathway, estrogen binding, and regulation of cell proliferation. Our strategy allowed the identification of putative casual mutations associated with muscle growth.

## BACKGROUND

Chicken is an important source of protein for human nutrition and a model system in growth and developmental biology (Ellegren 2005). The complete genome sequence of a Red Jungle Fowl female (*Gallus gallus gallus*), that is considered the ancestor of domestic chicken (*G. g. domesticus*) (Abplanalp 1992; Cassoli 2007; Dodgson *et al*. 2011), was completed in 2004 (Hillier *et al*. 2004) and opened the opportunity to explore the molecular control of complex phenotypes such as growth and muscle deposition among other traits.

High throughput sequencing of several chicken lines allowed the identification of millions of single nucleotide polymorphisms (SNPs) in the chicken genome (Rubin *et al*. 2010; Boschiero *et al*. 2018) and develop high density SNP panels (Kranis *et al*. 2013). SNPs are the most common and frequent DNA variant, with approximately 5 SNPs per kilobase (kb) in chicken (Rubin *et al*. 2010). When located in coding and regulatory regions of genes, they may affect traits of economic interest in animal models and livestock species (Roux *et al*. 2014).

High-density SNP panels were used in genome wide association studies (GWAS) to identify genomic regions associated with quantitative traits such as body weight (Gu *et al*. 2011a), fatness traits (Sun *et al*. 2013), breast and leg muscle weight, wing weight (Xie *et al*. 2012), carcass and eviscerated weight (Liu *et al*. 2013).

GWAS relies on the linkage disequilibrium of the genetic variant present in the SNP panel and the casual mutation, so further studies are necessary to identify the mutation responsible for the phenotype of interest. In an attempt to solve this issue, statistical evidence, such as association studies combined with functional annotations of genes and genetic variants, is important to determine he causal mutation (Spain and Barrett 2015). A SNP that occurs in coding regions can be classified as missense when the coded amino acid is changed, or synonymous, when the coded amino acid remains the same. Thereby, missense SNPs can be predicted as deleterious or tolerated by SIFT tool [Sorting Intolerant From Tolerant, (Ng and Henikoff 2003)]. Changes at well-conserved positions tend to be deleterious due to the assumption that important amino acids will be conserved in the protein family. (Ng and Henikoff 2002, 2003). When a SNP is predicted as a deleterious mutation, it means that the change of amino acids probably affects the protein structure and function, and consequently, may potentially alter the phenotype.

Using missense SNPs, some previous studies identified associations with body weight at hatch, semi-eviscerated carcass weight, eviscerated carcass weight, leg muscle weight and carcass weight (Wang *et al*. 2015), abdominal fat weight, body weight at different ages and body size traits (Han *et al*. 2012). However, there are no studies in the literature using predicted deleterious SNPs for association studies to identify casual mutations. Therefore, in this study we used previously developed whole genome sequence and GWAS information to identified predicted deleterious SNPs in QTL regions. Furthermore, we tested the association of these SNP with carcass traits in order to identify potential causal mutations in broilers.

## METHODS AND MATERIALS

### Ethics statement

In this study, all experimental protocols that used animals were performed in agreement with the resolution number 010/2012 approved by the Embrapa Swine and Poultry Ethics Committee on Animal Utilization to ensure compliance with international guidelines for animal welfare.

### Experimental Population

The TT reference population used for this study was generated from an Embrapa broiler line called TT. The TT line has been under selection since 1992, for many generations and several traits, with the goals to increase body weight and carcass yield, improve viability, fertility, hatchability, feed conversion, and reduce abdominal fat (Rosário *et al*. 2009). The TT Reference Population is an expansion of the TT line and was developed from crossing 20 males and 92 females (1:5) in five hatches, yielding approximately 1,500 chickens (Cruz et al. 2015; Marchesi et al. 2017). From this population we selected 237 offspring and 37 parental chickens (12 males and 25 females) for target sequencing. The offspring were selected based on the following criteria: (1) descendent of one of the 14 parental males that we have the whole genome sequencing data; (2) from families that have between 5 to 7 animals; (3) hatched on the first three incubations.

### Phenotype measurement

Body weight at 42 days of age (BW42) was measured six hours after fasting. Blood samples were collected for DNA extraction during the bleeding. After bleeding feathers were removed mechanically following a hot water bath (60°C for 45 s). The carcass cuts as breast weight (BTW), thigh weight (THW), drumstick weight (DRW) and abdominal fat weight (ABFW) were individually measured in grams. Drumstick yield (DR%), abdominal fat yield (ABF%), thigh yield (TH%) and breast weight (BT%) were estimated as a percentage of live body weight at 42 days of age (BW42). More details about the slaughter and phenotypes measurements are available at (Venturini *et al*. 2014; Cruz *et al*. 2015).

### SNPs selection and custom amplicon design

Predicted deleterious SNPs were selected from whole genome re-sequence data previously generated from 14 of the 37 parental animals of the population used in this study (TT Reference population). Sequences were generated with an Illumina HiSeq and SNP identified using SAMtools v.1.2 software (Li *et al*. 2009). Further details about library preparation, sequencing and filtering are available in Moreira *et al*. (2015) and Boschiero *et al*. (2018). SNP functional annotation was performed using VEP (Variant Effect Predictor, McLaren *et al*. 2016) and deleterious prediction was based on SIFT score prediction (Sorting Intolerant From Tolerant, Ng and Henikoff 2003). All the SNPs identified are available at EVA-EMBL database (https://www.ebi.ac.uk/ena/data/view/PRJEB25004).

In addition, a GWAS was performed by Moreira *et al*. (unpublished data) in the same meat-type chicken population (TT), in which some QTLs associated with BTW, BT%, THW, TH%, DRW, DR%, ABFW and ABF% traits were identified using high-density SNP chip (600K) data.

For SNPs selection, we overlapped all predicted deleterious SNPs identified in parental animals with the genomic windows identified in GWAS analysis that explained greater than 0.53 percent of the additive genetic variance associated with the studied traits (Supplemental Material, Table S1). Afterwards, the overlapped SNPs were analyzed with Tagger tool in Haploview software (Barrett *et al*. 2005) based on the linkage disequilibrium. In existence of adjacent SNPs with r^2 >^ 0.3, just one SNP remained to avoid the selection of markers in linkage disequilibrium and consequently, markers accounting for the same effect.

We defined 150 bp around each predicted deleterious SNP as a target region, with the variant located in the middle of the region. These regions were selected for target sequencing, and the amplicons designed were performed using DesignStudio online platform (Illumina Technology).

### Target sequencing

Genomic DNA was extracted using PureLink® Genomic DNA kit (Invitrogen, Carlsbad, CA, USA) and quantified using Qubit® 2.0 Fluorometer (Thermo Fisher Scientific, Waltham, MA, USA). DNA integrity was evaluated in 1% agarose gel. Library preparation was performed according to Truseq® Custom Amplicon Low Input Kit Reference Guide (Illumina Technology). Libraries were quantified with quantitative real time PCR, using KAPA® Library Quantification kit (KAPA Biosystem) and size estimated using either Bioanalyzer® (Agilent Technologies) or Fragment Analyzer (Advanced Analytical Technologies). Paired-end sequencing with a read length of 150 bp was performed on a MiniSeq™ (Illumina Technology).

### Sequencing data analyses, variant calling and functional annotation

The SNP calling was conducted for the 282 chickens (offspring and parental generations). Raw sequencing data were aligned against the chicken reference genome *Gallus_gallus*5.0 (NCBI) with BWA v.0.7.15 program, using BWA-MEM algorithm. For the SNP calling, we used SAMtools v.1.3.1 program (Li *et al*. 2009), with *mpileup* option (Li 2011), and mapping and base qualities (Phred) ≥ 20. The variant calling was performed with all 282 animals together. After the initial variant identification, the following filtering options were applied: INDEL removal, minor allele frequency (MAF) ≥ 0.05, SNP call rate ≥ 0.7, biallelic locus, sequencing depth ≥ 15 and Phred score quality ≥ 40.

After variant calling and filtration, the remainig SNPs were annotated using VEP tool version 91 (McLaren *et al*. 2016) available on Ensemble v. 91 website (Zerbino *et al*. 2018), and the SIFT score was predicted.

### Linkage disequilibrium analysis

Linkage disequilibrium analysis was conducted to indentify SNPs in strong linkage and avoid testing SNPs that capture the same effects. Predicted deleterious SNPs were visualized in Haploview program (Barrett *et al*. 2005) and adjacent SNPs with r^2^ > 0.8 were considered having a strong LD, and consequently one of them was excluded from association analyses. Haploview was also used to determine if the SNPs were in Hardy-Weinberg equilibrium (HWE).

### Association analysis

For association analysis, Proc Mixed Procedure was used on SAS 9.4 Studio online platform (Statistical Analysis System Institute Inc., Cary, NC). Association analysis was performed with all 20 predicted deleterious SNPs together for each carcass trait, and because of that correction for multiple tests was not necessary. The model used was:

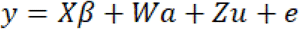

Where *y* is the vector of observations for the measured phenotype; *X* is the incidence matrix relating the fixed effects to **y**; *β* is the vector of fixed effects, which included sex and incubation; *W* is the genotype matrix (coded as 0, 1 and 2; 0 and 2 for homozygous and 1 for heterozygous) for all 20 deleterious SNPs and *a* is the vector of SNPs fixed effects. *Z* is the incidence matrix relating *u* to *y*; *u* is the vector of the family random effect; and *e* is the vector of residual effects. For the weight traits, BW42 was used as a covariate. Association was considered significant at p-value < 0.05 for the F test.

Orthogonal contrasts were used to compare the mean performance of one homozygote against another and to estimate additive and allelic substitution effects. Similarly, dominance effect was estimated through the orthogonal contrast of the mean performance of the heterozygote against the mean performance of both homozygotes. These analyses were performed under the same linear model detailed above with Proc Mixed Procedure on SAS 9.4 Studio online platform, considering the SNPs that presented the three genotypes (0, 1 and 2). Estimates and contrasts were set based on the methodology defined by Falconer and Mackay (1996). Effects were considered significant for p-value < 0.05 in the F test.

### Data availability

Data and reagents are available upon request. Table S1 contain the characterization of genomic windows identified in genome wide association analysis.

## RESULTS

### Phenotype measures

The summary statistics for BW42, THW, TH%, ABFW, ABF%, DRW, DR%, BTW and BT% are given in Table 1.

**Table 1.** Number of animals (N), mean, standard deviation (SD), minimum and maximum values for body weight at 42 days of age (BW42), thigh weight (THW), thigh yield (TH%), abdominal fat weight (ABFW), abdominal fat yield (ABF%), drumstick weight (DRW), drumstick yield (DR%), breast weight (BTW) and breast yield (BT%).

| Phenotype | N | Mean | SD | Minimum | Maximum |
| --- | --- | --- | --- | --- | --- |
| **BW42 (g)** | 237 | 2219.75 | 254.87 | 1310.00 | 2816.00 |
| **THW (g)** | 237 | 203.75 | 30.27 | 110.80 | 277.80 |
| **TH%** | 237 | 9.16 | 0.63 | 7.26 | 11.64 |
| **ABFW (g)** | 237 | 50.93 | 14.63 | 19.00 | 91.00 |
| **ABF%** | 237 | 2.29 | 0.61 | 1.01 | 4.25 |
| **DRW (g)** | 237 | 30.01 | 44.63 | 161.60 | 419.20 |
| **DR%** | 237 | 13.80 | 0.89 | 11.81 | 16.35 |
| **BTW (g)** | 237 | 495.39 | 61.93 | 260.00 | 660.00 |
| **BT%** | 237 | 22.32 | 1.31 | 18.28 | 26.51 |

### SNP selection and amplicon design

The whole genome re-sequencing from 14 parental chickens of the studied meat-type population identified approximately 11 million SNPs across the genome, and after functional annotation 4,708 of them were predicted as deleterious SNPs. As the result of the overlap between these 4,708 SNPs and the six selected genomic windows identified in GWAS, 89 predicted deleterious SNPs were kept for the further analysis. After linkage disequilibrium analysis (r^2^>0.3), 20 uncorrelated SNPs remained for the amplicon design using DesignStudio online platform. The final amplicon panel had 99% of coverage.

### Sequencing and variant calling

Libraries sequencing from MiniSeq produced an average number of raw reads of 298,791 per sample. The average of overall mapping rate of the raw reads against the *Gallus_gallus*5.0 (NCBI) genome assembly was 99.74%. After variant calling, 1,957 variants (including SNPs and INDELs) were initially detected, and 195 SNPs remained after filtration. The average depth of the remaining SNPs was 7,961 reads.

Functional annotation was performed for the 195 SNPs. As shown in Table 2, 29 SNPs were annotated as novel variants. From the 195 SNPs, 26% were in intronic regions, 20% were classified as missense and 14% were synonymous variants.

**Table 2.**
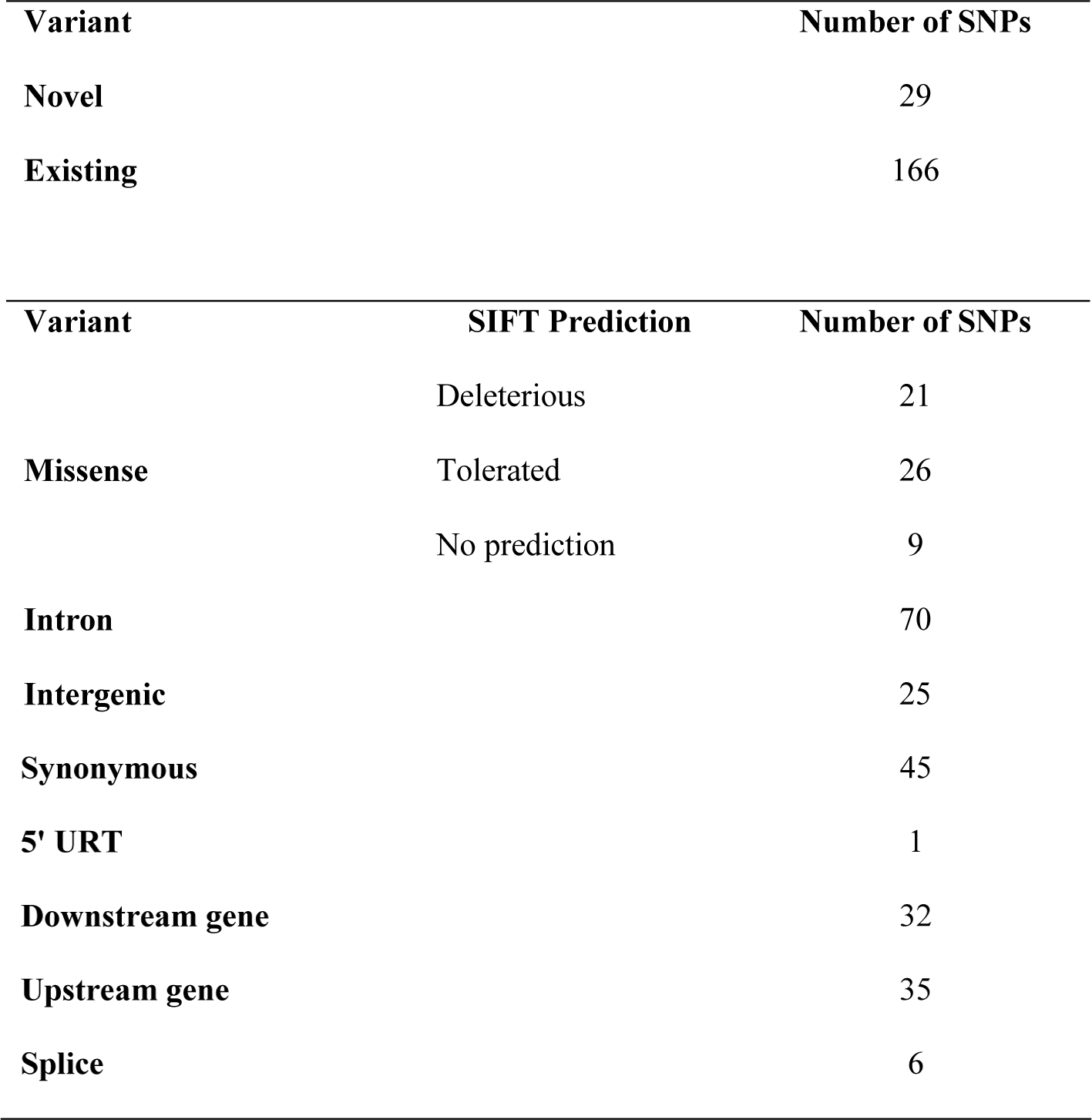
Number of novel and existing variants, and classification of functional annotation of single nucleotide polymorphisms (SNPs) performed with Variant Effect Predictor (VEP) online platform.

| Variant | Number of SNPs |
| --- | --- |
| Novel | 29 |
| Existing | 166 |

| Variant | SIFT Prediction | Number of SNPs |
| --- | --- | --- |
| Missense | Deleterious | 21 |
|  | Tolerated | 26 |
|  | No prediction | 9 |
| Intron |  | 70 |
| Intergenic |  | 25 |
| Synonymous | 45 |
| 5' URT | 1 |
| Downstream gene | 32 |
| Upstream gene | 35 |
| Splice | 6 |

As already mentioned, missense SNPs can be predicted as deleterious or tolerated based on its SIFT score (Ng and Henikoff 2003). In our study, from the 56 missense SNPs identified, 26 were classified as tolerated, 21 as deleterious, and 9 had no prediction (Table 2).

### Linkage disequilibrium verification

As presented in Figure 1, only one region of adjacent SNPs had *r*^*2*^ > 0.3 (*r*^*2*^ = 1.0, black square – chr26:3747344 and chr26: 3747346). Thus, chr26:3747344 was excluded, leaving 20 predicted deleterious SNPs for further analysis.

**Figure 1.**
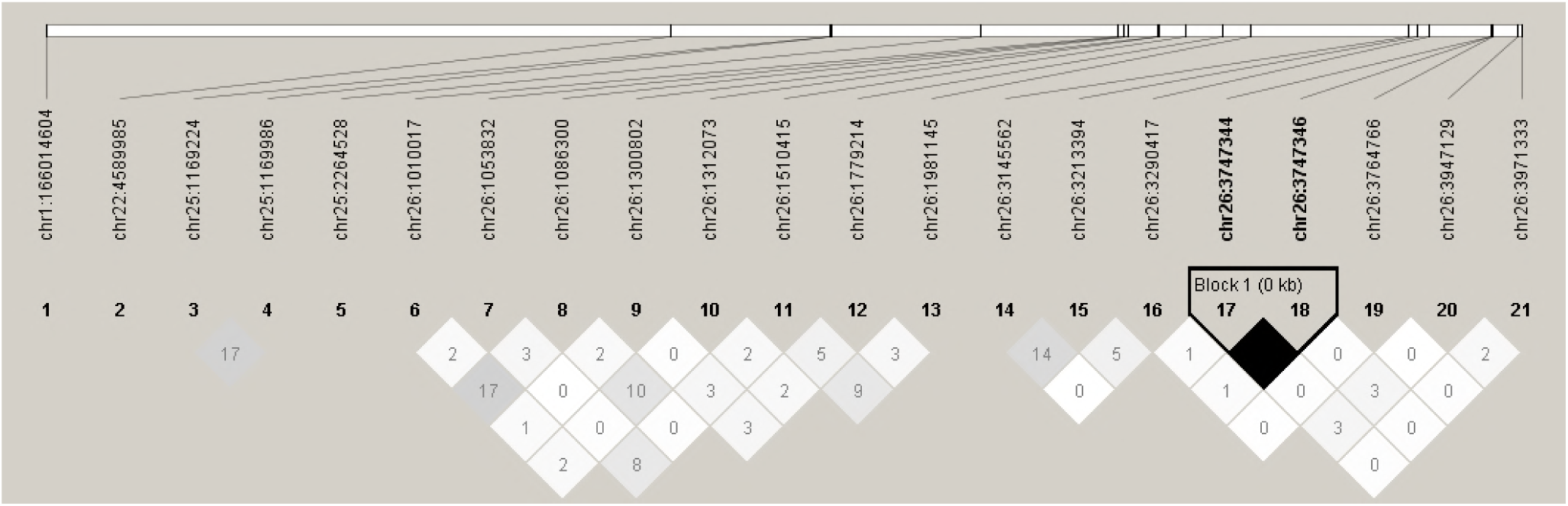
Linkage disequilibrium plot of the 21 predicted deleterious SNPs after filtering steps. The r^2^ value is presented within each square. The gradient color also represents r^2^ values, white is 0 and black is 1.

Detailed information of the 20 predicted deleterious SNPs (genome position, SNP ID, located gene, alleles and genotypes frequencies, HWE test and SIFT score) is presented in Table 3. Two SNPs did not have *rs* ID, and four genes were considered as novel genes, therefore, the ensemble gene ID was also presented. Seven SNPs did not have any animal genotyped with the alternative homozygous, and five SNPs were significant for HWE test.

**Table 3.**
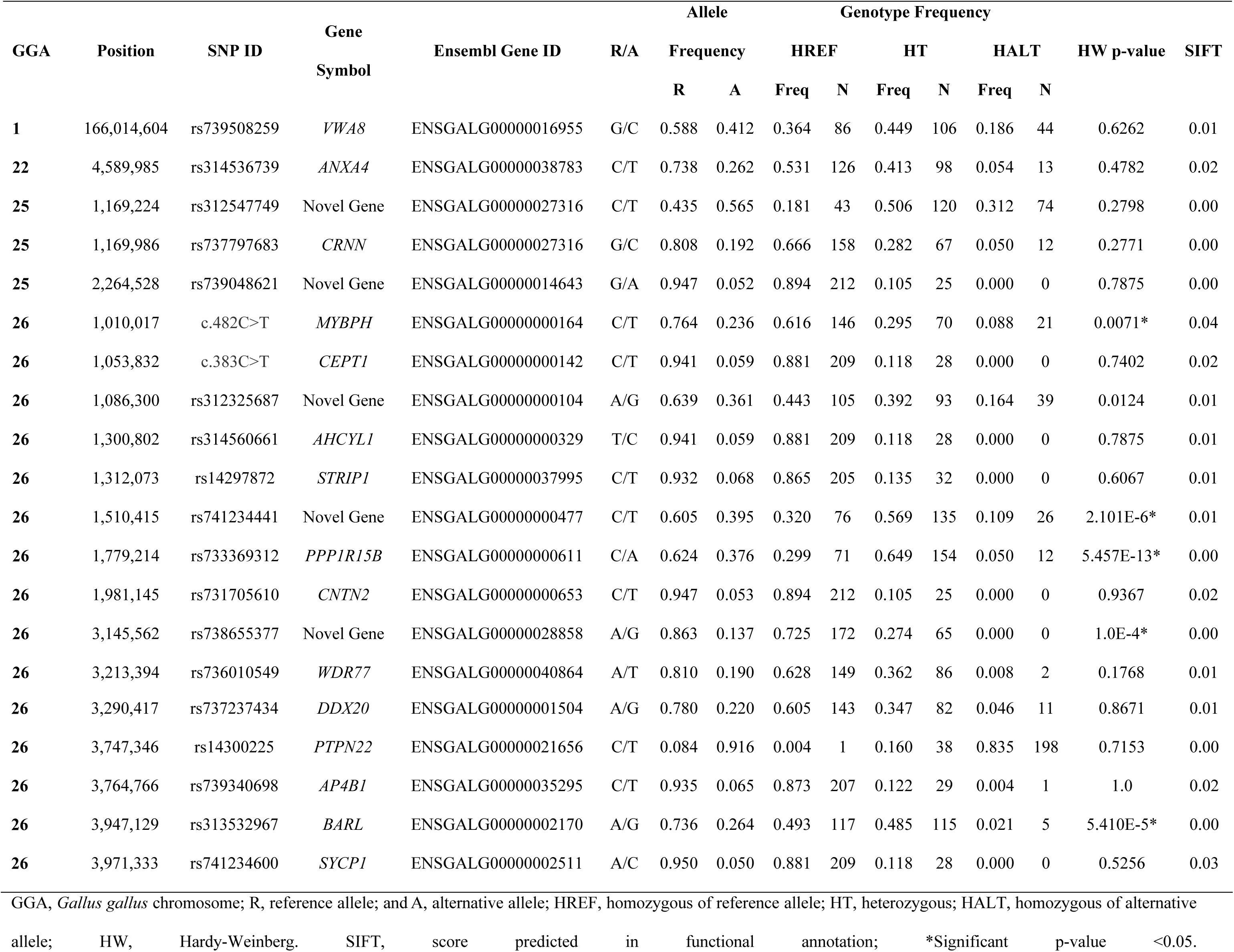
Deleterious SNPs selected for the association analyses with carcass traits.

### Association analysis, additive and dominance effects

Of the 20 predicted deleterious SNPs studied, six were significantly associated (p- value<0.05) with at least one carcass trait. Three SNPs (rs737797683, rs313532967 and rs741234600) were associated with breast traits; another three (rs739508259, rs312325687 and rs741234600) were associated with thigh traits; and one SNP (rs736010549) was associated with drumstick weight. No SNP was associated with abdominal fat traits. Detailed results for all association tests (p-values) are presented in Table 4.

**Table 4.**
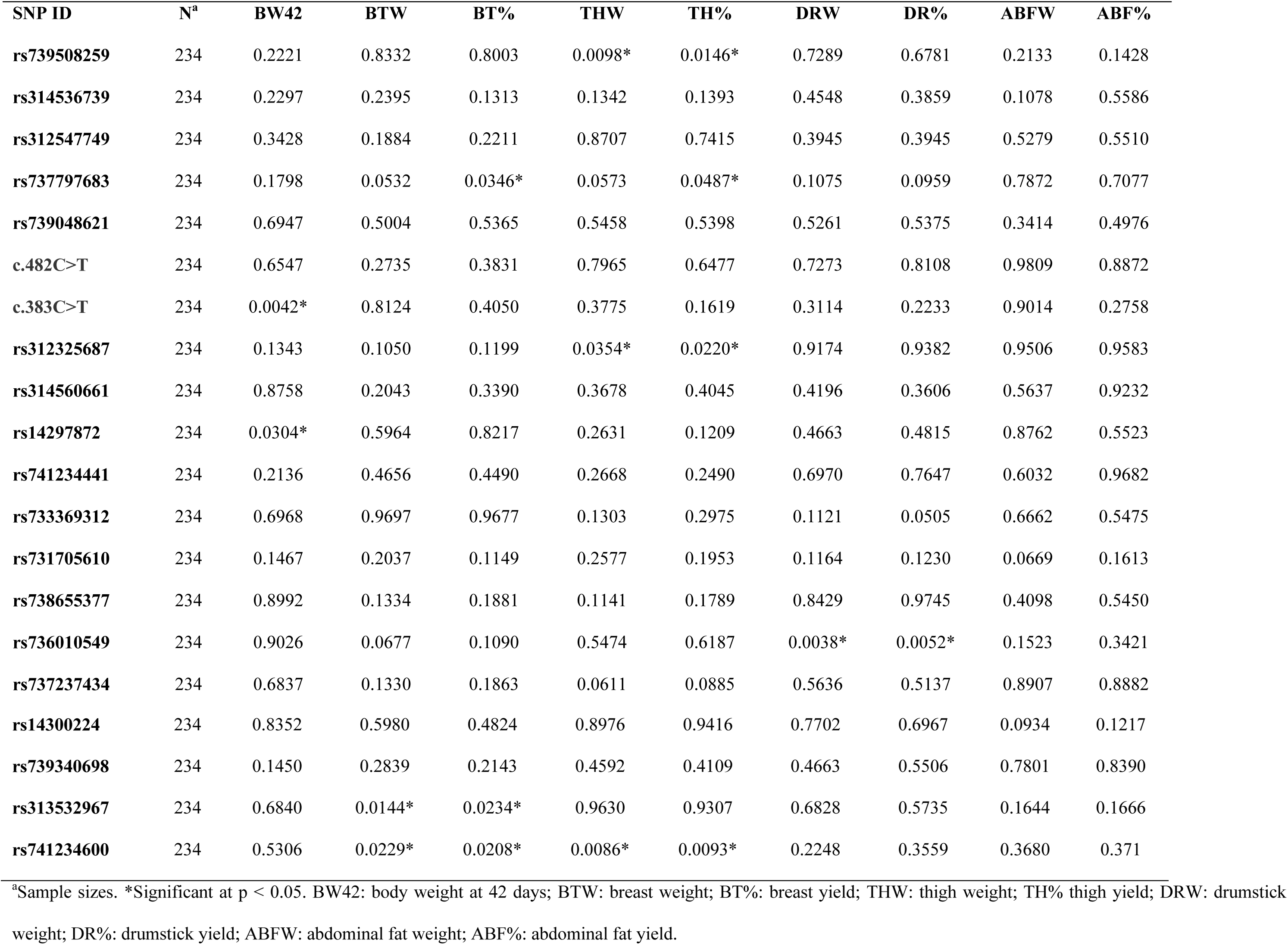
SNPs association analyses results for carcass traits.

Additive and dominance effects were estimated only for associated SNPs that presented the three genotypes, consequently rs741234600 was not considered in these analyses. We deemed significant effects with p-value <0.05. Allele substitution effect test was performed only for significant additive effects (Table 5). Additive and allele substitution effects for rs739508259 and rs312325687 associated with THW and TH% were significant. Additive and dominance effects for rs736010549 associated with DRW and DR% were significant, but not for allele substitution test.

**Table 5.** P-values, estimates and standard error for additive, dominance and allele substitution effects for SNPs with respective associated traits in broilers.

| SNP ID | Association |  | Additive Effect |  | Dominance Effect |  | Allele Substitution Effect |  |
| --- | --- | --- | --- | --- | --- | --- | --- | --- |
|  | Trait | p-value | P | E (se) | P | E (se) | P | E (se) |
| rs739508259 | THW | 0.0098** | 0.0029* | -5.7594 (1.9068) | 0.7616 | -0.7945 (2.6155) | 0.0062* | 5.1219 (1.8496) |
|  | TH% | 0.0146* | 0.0042* | -0.2479 (0.08523) | 0.6723 | -0.04971 (0.1174) | 0.0064* | 0.2268 (0.08213) |
| rs737797683 | BT% | 0.0346* | 0.0104* | -0.4857 (0.1880) | 0.2308 | -0.2553 (0.2124) | 0.0506 | 0.3041 (0.1547) |
|  | TH% | 0.0487* | 0.1966 | -0.1902 (0.1469) | 0.0144* | -0.4123 (0.1673) | - | - |
| rs312325687 | THW | 0.0354* | 0.0124* | 6.6692 (2.6307) | 0.9484 | -0.1913 (2.9505) | 0.0126* | -6.5551 (2.5853) |
|  | TH% | 0.0220* | 0.0069* | 0.3185 (0.1160) | 0.9387 | 0.01018 (0.1323) | 0.0063* | -0.3148 (0.1130) |
| rs736010549 | DRW | 0.0038* | 0.0009* | -13.8438 (4.1322) | 0.0016* | -13.8035 (4.3241) | 0.2394 | 2.3175 (1.9641) |
|  | DR% | 0.0052* | 0.0016* | -0.5862 (0.1836) | 0.0017* | -0.6107 (0.1919) | 0.3520 | 0.08240 (0.08834) |
| rs313532967 | BTW | 0.0144* | 0.6572 | -2.4332 (5.4758) | 0.2326 | 6.9216 (5.7825) | - | - |
|  | BT% | 0.0234* | 0.6708 | -0.1085 (0.2550) | 0.2634 | 0.3015 (0.2689) | - | - |
| rs741234600 | BTW | 0.0229* | - | - | - | - | 0.0602 | 9.4399 (4.9996) |
|  | BT% | 0.0208* | - | - | - | - | 0.0491* | 0.4620 (0.2335) |
|  | THW | 0.0086* | - | - | - | - | 0.0136* | 10.0902 (4.0577) |
|  | TH% | 0.0093* | - | - | - | - | 0.0121* | 0.4598 (0.1819) |

THW: thigh weight; TH% thigh yield; DRW: drumstick weight; DR%: drumstick yield; BTW: breast weight; BT%: breast yield. P, E and Se are p-values, estimates and standard errors for the respective analysis. \*Significant at p < 0.05.

## DISCUSSION

The identification of genetic markers associated with carcass weight and yield traits has been the focus of several studies due to the economic importance of these traits in broiler production. With the main goal of finding putative causal mutations for carcass traits, this study selected predicted deleterious SNPs present in QTLs regions to be evaluated as potential causal mutations in our TT Reference Population.

The SNP rs739508259 is located in the von willebrand factor A domain containing 8 (*VWA8*) gene and within the GGA-1 at 166 Mb genomic window identified in the GWAS analysis. This region was associated with DRW and DR%, explaining 3.20 and 2.79 of the additive genetic variance, respectively. This SNP is a G>C nucleotide change with minor allele (C) frequency of 0.42. The nucleotide change causes the amino acid substitution of glutamine to histidine. The rs739508259 was associated with THW and TH% and had significant additive and allele substitution effects for both traits. On average, for each C allele in the animal’s genotype, an increase of 5.12g was observed for THW and 0.24% for TH%, compared to the GG genotype. The window on GGA-1 at 166 Mb also explained 0.20% and 0.14% of the additive genetic variance for THW and TH%, respectively. However, these proportions were not enough to be considered significant (Additional File 1). Furthermore, it is interesting to observe that SNP rs739508259 was not significantly associated with DRW and DR% and this may be because we used a subset of 237 animals from the 1,408 animals used in GWAS analysis or because the low number of animals used (237) in the association analysis. Similar findings were also noted for other associated SNPs.

In mice *VWA8* gene is highly expressed in skeletal muscle, has ATPase domains, mitochondrial targeting sequences and is a mitochondrial protein (Luo *et al*. 2017). More studies are necessary to relate this gene with muscle growth in chickens.

The predicted deleterious SNP rs736010549 is located in the WD repeat domain 77 (*WDR77*) gene. Furthermore, it is located within the GGA-26 at 3 Mb genomic window identified in GWAS analysis, and this region was associated with breast weight (BRW) and breast yield (BR%), representing 0.53 and 0.86 of the additive genetic variance, respectively (Additional File 1). This polymorphism results in an amino acid change from serine to cysteine (A/T allele substitution), with the minor allele (T) frequency of 0.19. This SNP was associated with drumstick weight (DRW) and drumstick yield (DR%). It was also significant for additive and dominance effects tests for both traits. On average, animals with TT genotype had 13.8g more of DRW and 0.58 % more of DR% than AA animals. This SNP presented complete dominance for both traits.

The WD repeat domain 77 (*WDR77*) gene belongs to the WD repeat proteins that is characterized by multiple protein interaction capacity (Friesen *et al*. 2001). The protein p44 (also named as methylosome protein 50, MEP50) coded by *WDR77* and is an androgen receptor (AR) coactivator by multiprotein complex formation (Hosohata *et al*. 2003). In humans, p44 was associated with inhibition of prostate cancer cell growth as coactivator of AR (Zhou *et al*. 2006; Gu *et al*. 2011b) and with breast cancer growth mediated through estrogen and its receptor (Peng *et al*. 2010).

Several studies showed the inhibitory action of androgenic steroids in chicken growth (Fennell *et al*. 1990; Fennell and Scanes 1992; Esquivel-Hernandez *et al*. 2016), which is a possible consequence of the androgen receptor or estrogen receptor aromatization (Fennel *et al*. 1996; Callewaert *et al*. 2010). Kong *et al*. (2017) studied different expressed (DE) genes in a selected and unselected broiler breeds, and among their results, they suggested that inhibited AR was predicted to be an effective regulatory factor for DE genes in selected breed, corroborating previously cited studies. Our results suggest that SNP rs736010549 alter the conformation of p44, decreasing the AR activation and so contributing to growth in chickens.

The predicted deleterious rs313532967 is located in the bile acid receptor-like (*BARL*) gene and within the GGA-26 at 3 Mb genomic window identified in GWAS analysis. This region was associated with BTW and BT%, representing 0.53 and 0.86 of the additive genetic variance respectively. This SNP is an A>G change, resulting in the amino acid change of asparagine to serine, and the minor allele (G) frequency is 0.264. The HWE test was significative, and this may be due to our finite population, or indicating that this locus may be under selection or inbreeding. This SNP was associated with BTW and BT% and did not have additive or dominant effects. Only five animals had GG genotype and this could help explain the lack of additive and dominant effects.

The *BARL* gene have a DNA-binding domain of Farnesoid X receptor (FXR) family. This domain in humans was intensively studied and when it is activated by bile acids it can regulate bile acids synthesis, conjugation and transport, consequently impacting in lipid and glucose metabolism (Claudel *et al*. 2005; Preidis *et al*. 2017). When bile acids are released in the ileum, its induces the synthesis of fibroblast growth factor (FGF-19) which stimulates hepatic protein and glycogen synthesis (Kir *et al*. 2011). In an interesting work in broilers, Lai *et al*. (2018) demonstrated that dietary supplementation of swine bile acids for broiler chickens influences their growth performance and carcass characteristics as reduction of abdominal fat, increase of carcass weight, eviscerated weight and leg weight. Therefore, our study indicates that *BARL* gene can be involved in growth and carcass development in chicken.

The SNP rs741234600 is located in synaptonemal complex protein 1 (*SYCP1*) gene and within the GGA-26 at 3 Mb genomic window identified in GWAS analysis. This region was associated with BTW and BT%, representing 0.53 and 0.86 of the additive genetic variance respectively. In our study, this SNP was associated with BTW, BT%, THW and TH%, and is an A>C nucleotide change, resulting in the amino acid substitution of lysine to threonine, and the minor allele (C) frequency is 0.05. As the CC genotype (alternative homozygous) was not present in the evaluated animals, only allele substitution effect was performed. For each allele A, the animals had 0.46% more BT%, 10.09 g more THW and 0.45% more TH%.

The SNP rs737797683 is located in the cornulin (*CRNN*) gene, within the GGA-25 at 1 Mb genomic window identified in GWAS analysis. This region was associated with BT%, BR% and ABF% representing 0.24, 0.24 and 0.23 of the additive genetic variance respectively. This SNP was associated with BT% and TH%. This SNP is an G>C change, resulting in the amino acid change of aspartic acid to histidine, and the minor allele (C) frequency is 0.19. This polymorphism was significant for additive effect for BT% trait, significant for dominance effect for TH%.

The SNP rs312325687 is located in the cryptochrome 4 (*CRY4*) gene and within the GGA-26 at 1 Mb genomic window identified in GWAS analysis, previously associated with ABFW and ABF%, representing 1.06 and 0.54 of the additive genetic variance respectively. This SNP was associated with THW and TH% and is an A>G change, resulting in the amino acid change of aspartic acid to glycine, being the minor allele (G) frequency equal to 0.36. This polymorphism had significant additive and allele substitution effects for both traits.

*SYCP1, CRNN, CRY4* were not selected as candidate genes for muscle growth or carcass development in this study because there is no information available in the literature to support this. More studies with these genes are necessary to understand their relationship with carcass and muscle growth.

It is pertinent to note that although rs738655377 was not significantly associated with any of the phenotypes tested, none of the animal were homozygous for the alternative allele, and the HWE test was significant. This observation provides evidence for a lethal polymorphism when in homozygosity. This variant is within a novel gene (ENSGALG00000028858) that has gene ontology terms related to oxidoreductase activity.

In conclusion, our study identified 20 predicted deleterious SNPs in different QTLs associated with carcass traits and succeeded in associating six of them with phenotypes related to muscle growth. Three predicted deleterious SNPs associated were located in genes that we consider candidate genes for carcass and muscle weight, and development. The main limitation of our study is that it is difficult to determine if the identified mutations are the causative mutation or are in linkage disequilibrium with the real causal mutation.

## ACKNOWLEDGMENTS

The authors are grateful for the support from São Paulo Research Foundation (FAPESP) thematic grant 14/08704-0 and Brazilian Agricultural Research Corporation – Embrapa (project number 01.11.07.002.04.02). The TT Reference Population was subsidized by the National Council of Scientific and Technological Development (CNPq) grant number 481755/2007-1 from the Brazilian Government. C. Boschiero received a fellowship from the program Science Without Borders - CNPq, grant 370620/2013-5. P.A. Trevisoli received fellowship from FAPESP grant 2016/13589-0. L.L. Coutinho is recipient of productivity fellowship from CNPq. The authors would like to acknowledge the collaborative efforts among EMBRAPA Suínos e Aves and University of São Paulo.

---

**Table S1.** Characterization of genomic windows identified in genome wide association analysis for the studied traits.

| **Trait** | **GGA_Mb<sup>a</sup>** | **Start – end positions**<br>**(Chr_SNP)** | **SNPs<sup>b</sup>** | **%GV<sup>c</sup>** | **PPA**<br>**(p>0)<sup>d</sup>** |
| --- | --- | --- | --- | --- | --- |
| **DRW** | 1_166 | 1_166000511 - 1_166999195 | 390 | 3.20 | 0.92 |
|  | 26_3 | 26_3000141 - 26_3998650 | 998 | 0.22 | 0.94 |
| **DR%** | 1_166 | 1_166000511 - 1_166999195 | 390 | 2.79 | 0.93 |
|  | 26_3 | 26_3000141 - 26_3998650 | 998 | 0.25 | 0.95 |
| **THW** | 1_166 | 1_166000511 - 1_166999195 | 390 | 0.20 | 0.71 |
|  | 22_4 | 22_4000760 - 22_4676714 | 1035 | 0.54 | 0.95 |
|  | 26_1 | 26_1002598 - 26_1999851 | 662 | 0.11 | 0.84 |
| **TH%** | 1_166 | 1_166000511 - 1_166999195 | 390 | 0.14 | 0.71 |
|  | 22_4 | 22_4000760 - 22_4676714 | 1035 | 0.57 | 0.97 |
|  | 25_1 | 25_1000996 - 25_1982441 | 691 | 0.11 | 0.85 |
|  | 26_1 | 26_1002598 - 26_1999851 | 662 | 0.14 | 0.85 |
| **BRW** | 25_1 | 25_1000996 - 25_1982441 | 691 | 0.24 | 0.88 |
|  | 25_2 | 25_2001192 - 25_2887176 | 512 | 0.81 | 0.88 |
|  | 26_3 | 26_3000141 - 26_3998650 | 998 | 0.53 | 0.96 |
| **BR%** | 25_1 | 25_1000996 - 25_1982441 | 691 | 0.24 | 0.90 |
|  | 25_2 | 25_2001192 - 25_2887176 | 512 | 0.6 | 0.84 |
|  | 26_3 | 26_3000141 - 26_3998650 | 998 | 0.86 | 0.98 |
| **ABFW** | 26_1 | 26_1002598 - 26_1999851 | 662 | 1.06 | 0.95 |
| **ABF%** | 25_1 | 25_1000996 - 25_1982441 | 691 | 0.23 | 0.87 |
|  | 26_1 | 26_1002598 - 26_1999851 | 662 | 0.54 | 0.92 |

<sup>a</sup>Map position based on Gallus\_gallus-5.0 NCBI assembly; <sup>b</sup>Number of SNPs per region; <sup>c</sup>% of genetic variance explained by the window; <sup>d</sup>Posterior probability of association (PPA) as reported by Onteru et al. (2013). DRW: drumstick weight; DR%: drumstick yield; THW: thigh weight; TW%: thigh yield; BTW: breast weight; BT%: breast yield; ABFW: abdominal fat weight; ABF%: abdominal fat yield;

## LITERATURE CITED

Abplanalp, H., 1992 Inbred lines as genetic resources of chickens. Poult. Sci. Rev. 4: 29–39.

Barrett, J. C., B. Fry, J. Maller, and M. J. Daly, 2005 Haploview: analysis and visualization of LD and haplotype maps. Bioinformatics 21: 263–265.

Boschiero, C., G. C. M. Moreira, A. A. Gheyas, T. F. Godoy, G. Gasparin et al., 2018 Genome-wide characterization of genetic variants and putative regions under selection in meat and egg-type chicken lines. BMC Genomics 19: 83.

Callewaert, F., M. Sinnesael, E. Gielen, S. Boonen, and D. Vanderschueren, 2010 Skeletal sexual dimorphism: relative contribution of sex steroids, GH-IGF1, and mechanical loading. J. Endocrinol. 207: 127–134.

Cassoli, C. S. da S., 2007 Identificação e análise de etiquetas de seqüências expressas (ESTs) na hipófise e hipotálamo de Gallus gallus. Tese-Esc. Super. Agric. Luiz Queiroz.

Claudel, T., B. Staels, and F. Kuipers, 2005 The Farnesoid X receptor: a molecular link between bile acid and lipid and glucose metabolism. Arterioscler. Thromb. Vasc. Biol. 25: 2020–30.

Cruz, V. A. R. da, F. S. Schenkel, R. P. Savegnago, N. V. Grupioni, N. B. Stafuzza et al., 2015 Association of Apolipoprotein B and Adiponectin Receptor 1 Genes with Carcass, Bone Integrity and Performance Traits in a Paternal Broiler Line (R. Davoli, Ed.). PLoS One 10: e0136824.

Dodgson, J. B., M. E. Delany, and H. H. Cheng, 2011 Poultry genome sequences: progress and outstanding challenges. Cytogenet. Genome Res. 134: 19–26.

Ellegren, H., 2005 The avian genome uncovered. Trends Ecol. Evol. 20: 180–186.

Esquivel-Hernandez, Y., R. E. Ahumada-Cota, M. Attene-Ramos, C. Z. Alvarado, P. Castañeda-Serrano et al., 2016 Making things clear: Science-based reasons that chickens are not fed growth hormones. Trends Food Sci. Technol. 51: 106–110.

Falconer, D. S., and T. F. C. Mackay, 1996 Introduction to Quantitative Genetics. Pearson Education Limited.

Fennel, M. J., S. V. Radecki, J. A. Proudman, and C. G. Scanes, 1996 The Suppressive Effects of Testosterone on Growth in Young Chickens Appears to be Mediated via a Peripheral Androgen Receptor; Studies of the Anti-Androgen ICI 176,334. Poult. Sci. 75: 763–766.

Fennell, M. J., A. L. Johnson, and C. G. Scanes, 1990 Influence of androgens on plasma concentrations of growth hormone in growing castrated and intact chickens. Gen. Comp. Endocrinol. 77: 466–475.

Fennell, M. J., and C. G. Scanes, 1992 Inhibition of Growth in Chickens by Testosterone, 5 - Dihydrotestosterone, and 19-Nortestosterone. Poult. Sci. 71: 357–366.

Friesen, W. J., A. Wyce, S. Paushkin, L. Abel, J. Rappsilber et al., 2001 A Novel WD Repeat Protein Component of the Methylosome Binds Sm Proteins*.

Gu, X., C. Feng, L. Ma, C. Song, Y. Wang et al., 2011a Genome-wide association study of body weight in chicken F2 resource population. PLoS One 6: e21872.

Gu, Z., L. Zhou, S. Gao, and Z. Wang, 2011b Nuclear transport signals control cellular localization and function of androgen receptor cofactor p44/WDR77. PLoS One 6: e22395.

Han, R., Y. Wei, X. Kang, H. Chen, G. Sun et al., 2012 Novel SNPs in the PRDM16 gene and their associations with performance traits in chickens. Mol. Biol. Rep. 39: 3153–3160.

Hillier, L. W., W. Miller, E. Birney, W. Warren, R. C. Hardison et al., 2004 Sequence and comparative analysis of the chicken genome provide unique perspectives on vertebrate evolution. Nature 432: 695–716.

Hosohata, K., P. Li, Y. Hosohata, J. Qin, R. G. Roeder et al., 2003 Purification and Identification of a Novel Complex Which Is Involved in Androgen Receptor-Dependent Transcription. Mol. Cell. Biol. 23: 7019–7029.

Kir, S., S. A. Beddow, V. T. Samuel, P. Miller, S. F. Previs et al., 2011 FGF19 as a Postprandial, Insulin-Independent Activator of Hepatic Protein and Glycogen Synthesis. Science (80-.). 331: 1621–1624.

Kong, B.-W., N. Hudson, D. Seo, S. Lee, B. Khatri et al., 2017 RNA sequencing for global gene expression associated with muscle growth in a single male modern broiler line compared to a foundational Barred Plymouth Rock chicken line. BMC Genomics 18:.

Kranis, A., A. A. Gheyas, C. Boschiero, F. Turner, L. Yu et al., 2013 Development of a high density 600K SNP genotyping array for chicken. BMC Genomics 14: 59.

Lai, W., W. Huang, B. Dong, A. Cao, W. Zhang et al., 2018 Effects of dietary supplemental bile acids on performance, carcass characteristics, serum lipid metabolites and intestinal enzyme activities of broiler chickens. Poult. Sci. 97: 196–202.

Li, H., 2011 A statistical framework for SNP calling, mutation discovery, association mapping and population genetical parameter estimation from sequencing data. Bioinformatics 27: 2987–2993.

Li, H., B. Handsaker, A. Wysoker, T. Fennell, J. Ruan et al., 2009 The Sequence Alignment/Map format and SAMtools. Bioinformatics 25: 2078–2079.

Liu, R., Y. Sun, G. Zhao, F. Wang, D. Wu et al., 2013 Genome-Wide Association Study Identifies Loci and Candidate Genes for Body Composition and Meat Quality Traits in Beijing-You Chickens (B. A. White, Ed.). PLoS One 8: e61172.

Luo, M., A. E. Mengos, W. Ma, J. Finlayson, R. Z. Bustos et al., 2017 Characterization of the novel protein KIAA0564 (Von Willebrand Domain-containing Protein 8). Biochem. Biophys. Res. Commun. 487: 545–551.

Marchesi, J. A. P., M. E. Buzanskas, M. E. Cantão, A. M. G. Ibelli, J. O. Peixoto et al., 2017 Relationship of runs of homozygosity with adaptive and production traits in a paternal broiler line. animal 1–9.

McLaren, W., L. Gil, S. E. Hunt, H. S. Riat, G. R. S. Ritchie et al., 2016 The Ensembl Variant Effect Predictor. Genome Biol. 17: 122.

Moreira, G. C. M., T. F. Godoy, C. Boschiero, A. Gheyas, G. Gasparin et al., 2015 Variant discovery in a QTL region on chromosome 3 associated with fatness in chickens. Anim. Genet. 46: 141–147.

Ng, P. C., and S. Henikoff, 2002 Accounting for human polymorphisms predicted to affect protein function. Genome Res. 12: 436–46.

Ng, P. C., and S. Henikoff, 2003 SIFT: Predicting amino acid changes that affect protein function. Nucleic Acids Res. 31: 3812–3814.

Peng, Y., Y. Li, L. L. Gellert, X. Zou, J. Wang et al., 2010 Androgen receptor coactivator p44/Mep50 in breast cancer growth and invasion. J. Cell. Mol. Med. 14: 2780–9.

Preidis, G. A., K. H. Kim, and D. D. Moore, 2017 Nutrient-sensing nuclear receptors PPARα and FXR control liver energy balance. J. Clin. Invest. 127: 1193–1201.

Rosário, M. F. do, M. C. Ledur, A. S. A. M. T. Moura, L. L. Coutinho, and A. A. F. Garcia, 2009 Genotypic characterization of microsatellite markers in broiler and layer selected chicken lines and their reciprocal F1s. Sci. Agric. 66: 150–158.

Roux, P.-F., M. Boutin, C. Désert, A. Djari, D. Esquerré et al., 2014 Re-sequencing data for refining candidate genes and polymorphisms in QTL regions affecting adiposity in chicken. PLoS One 9: e111299.

Rubin, C.-J., M. C. Zody, J. Eriksson, J. R. S. Meadows, E. Sherwood et al., 2010 Wholegenome resequencing reveals loci under selection during chicken domestication. Nature 464: 587–591.

Spain, S. L., and J. C. Barrett, 2015 Strategies for fine-mapping complex traits. Hum. Mol. Genet. 24Spain, S: R111–R119.

Sun, Y., G. Zhao, R. Liu, M. Zheng, Y. Hu et al., 2013 The identification of 14 new genes for meat quality traits in chicken using a genome-wide association study. BMC Genomics 14: 458.

Venturini, G. C., V. A. R. Cruz, J. O. Rosa, F. Baldi, L. El Faro et al., 2014 Genetic and phenotypic parameters of carcass and organ traits of broiler chickens. funpecrp.com.br Genet. Mol. Res. Genet. Mol. Res 13: 10294–10300.

Wang, Y. C., R. R. Jiang, X. T. Kang, Z. J. Li, R. L. Han et al., 2015 Identification of single nucleotide polymorphisms in the ASB15 gene and their associations with chicken growth and carcass traits. funpecrp.com.br Genet. Mol. Res. Genet. Mol. Res 14: 11377–11388.

Xie, L., C. Luo, C. Zhang, R. Zhang, J. Tang et al., 2012 Genome-Wide Association Study Identified a Narrow Chromosome 1 Region Associated with Chicken Growth Traits (Z. Liu, Ed.). PLoS One 7: e30910.

Zerbino, D. R., P. Achuthan, W. Akanni, M. R. Amode, D. Barrell et al., 2018 Ensembl 2018. Nucleic Acids Res. 46: D754–D761.

Zhou, L., H. Wu, P. Lee, and Z. Wang, 2006 Roles of the androgen receptor cofactor p44 in the growth of prostate epithelial cells. J. Mol. Endocrinol. 37: 283–300.

